# Prediction of disordered regions in proteins with recurrent Neural Networks and protein dynamics

**DOI:** 10.1101/2020.05.25.115253

**Authors:** Gabriele Orlando, Daniele Raimondi, Francesco Codice, Francesco Tabaro, Wim Vranken

## Abstract

The role of intrinsically disordered protein regions (IDRs) in cellular processes has become increasingly evident over the last years. These IDRs continue to challenge structural biology experiments because they lack a well-defined conformation, and bioinformatics approaches that accurately delineate disordered protein regions remain essential for their identification and further investigation. Typically, these predictors use only the protein amino acid sequence, without taking into account likely emergent properties that are sequence context dependent, such as protein backbone dynamics.

The DisoMine method predicts protein disorder with recurrent neural networks not directly from the amino acid sequence, but instead from more generic predictions of key biophysical properties, here protein dynamics, secondary structure and early folding. The tool is fast and requires only a single sequence, making it applicable for large-scale screening, including poorly studied and orphan proteins. DisoMine compares well to 10 state of the art predictors, also if these use evolutionary information.

DisoMine is freely available through an interactive webserver at http://bio2byte.com/disomine/

## 1 Introduction

The view of proteins as molecules with a stable and well-defined conformation has been, and continues to be, very useful to understand many of their characteristics, for example how an active site in an enzyme catalyses a reaction. However, a single rigid structure often fails to capture the full behavior of a protein. In the extreme case of intrinsically disordered regions (IDRs) or proteins (IDPs), the protein (or a region thereof) adopts a heterogeneous mix of conformations in fast exchange with each other. These proteins are therefore highly dynamic, with fast movements between many different conformational states.

Experimental studies, for example using NMR, have highlighted the importance of such non-globular and disordered proteins [1], which often have crucial functions with essential roles in cell life [2]. Recent studies [3] have also suggested that the portion of disordered proteins in the cell may be much larger than previously thought. However, this class of proteins is still not well studied and understood, with their highly dynamic and conformationally heterogeneous behavior challenging both structural biology methods and molecular dynamics (MD) approaches. Despite advances in these areas, especially in NMR [4] and in MD force field development [5, 6], bioinformatics tools that accurately predict protein disorder remain essential for both sequence and structure analysis. This is attested by the large number of IDR predictors that have been, and are being, developed [7]. The quality of such disorder predictions can have a large impact on the conclusions of computational and experimental studies [8] and, therefore, the development of faster and more accurate methods is essential for this rapidly progressing field of research.

The collection and curation of disorder annotation in public databases has been essential for the development of this field. Particularly important are the MobiDB and DisProt databases [9, 10]. MobiDB was developed in 2014 and is continuously updated, as it contains the integrated disorder predictions and annotations of several databases. DisProt, on the other hand, collects high quality, manually curated disorder annotations based on available experimental data. It was created in 2006 and is now the standard for research in the field. DisProt is especially focused on longer disordered regions.

In this paper we present DisoMine, a Neural Network (NN) based predictor of protein disorder regions that provides accurate predictions with respect to currently available tools. These performances were assessed on a recent publicly available and manually curated disorder dataset, which was previously used for a large scale disorder prediction benchmark [11]. The key novelty that sets our method apart is that disorder is not predicted directly from the amino acid sequence, but is instead computed from predicted properties of amino acid residues, such as their backbone dynamics [12]. The predictors for these biophysical properties are based on independent experimental datasets with quantitative information or well-delineated properties, unlike the order/disorder categories, which are difficult to capture. These independently calculated properties can then be assembled into feature vectors of continuous bounded values that effectively replace the amino acid code for machine learning methods, but which depend on the sequence context of the individual amino acid residues. This can significantly improve prediction results [13, 14], likely because the emergent properties are not obvious from the amino acid codes themselves, and are more useful after explicit inclusion. In the case of disorder prediction, such emergent properties can capture the underlying reasons of why a protein region tends to be disordered, and so help to make the disorder predictions more generally applicable. DisoMine has also already been successfully used in a pipeline for the identification of prion-like RNA-binding proteins that form liquid phase-separated condensates [15], defining the disorder content of the regions that typically constitute this class of proteins.

A web interface is publicly available at http://bio2byte.com/disomine/. It allows the user to obtain disorder predictions, as well as early folding residues and backbone dynamics estimated with the tools described in [14] and [12].

## 2 Methods

### 2.1 Datasets

DisoMine was trained on the DisProt dataset described from [16], which consists of 535 non-redundant proteins extracted from the DisProt database [10]. Every residue is manually annotated as “structured” or “disordered”, and these two states correspond to our prediction classes, with disordered residues being the positive cases. The protein lengths range from 32 to 7962 and residues, and the dataset contains a total of 236930 amino acids. Overall, 37% of the residues are annotated as disordered, 63% as structured (ordered).

For the validation of the method, we followed the procedure used for a recent and comprehensive disorder prediction benchmark [11]. The test set (*assessment dataset*) consists of proteins from the latest version of Disprot (version 7.0) that were not present in the previous releases. Similarly to our training set, this validation set contains proteins with manually annotated disordered regions, and comprises 268 proteins with no evolutionary correlation with the training set. Since DisoMine is both trained and tested on Disprot data, it is aimed at predicting typically longer disordered regions (the median disordered region length in the training set is 34 residues).

As an additional validation we used two additional datasets. The first one is a crystal structures-based dataset, obtained from [16]. The dataset is made of 3738 proteins, that consist in a total of 741906 residues, 47422 of which are annotated as disordered. Annotations are based on their presence (or absence) in the corresponding PDB structure, so encompassing especially flexible loop regions. The median length of the disordered regions is 8 residues. The second one is the one used for the CASP7 competition. It is made of 96 proteins, with a total of 18627 residues, 1189 of which annotated as disordered. The median disordered region length is again 8 residues.

### 2.2 Features

DisoMine’s input features consist of the predictions of the following bioinformatics tools: the PSIPRED secondary structure predictor (single sequence version) [17], the DynaMine protein backbone dynamics predictor [12], the EFoldMine early folding regions predictor and the DynaMine protein side-chain dynamics predictor described in [14]. All these tools compute their predictions using only the single target protein sequence as input. They do not use evolutionary information such as Multiple Sequence Alignments (MSAs) at any point, thus reducing DisoMine’s bias towards well studied proteins, which can be a significant problem in bioinformatics [18].

For each residue in each sequence, PSIPRED predicts Coil, Helix and Sheet propensities. We encode this information in a 3-dimensional vector for each residue. Similarly, backbone and sidechain dynamics as well as early folding propensity predictions result in another 3 values for each residue. We assemble these two vectors to obtain a 6-dimensional ‘biophysical’ vector describing each residue, and use these within a 7-residue window, with the central residue the one to be predicted. In our feature encoding scheme, a protein of length *L* is thus represented by a tensor with shape *L ×* 42.

### 2.3 The Neural Network model

DisoMine is implemented as a Recurrent Neural Network (RNN) using version 0.3.1 of `pytorch` [19]. The network is composed of two sub modules: the first one consists of a 2 layer bidirectional Gated Recurrent Unit (GRU) network [20] with 30 hidden dimensions and Rectified Linear Unit (ReLU) activations (Figure 1). The GRU architecture is a type of recurrent layer, meaning that it can take the sequential information of the protein sequence into consideration. Respect to other types of recurrent layers, such as Long Short Term Memory, it has fewer weights to tune since it does not parameterize the output gate. This is useful to keep the number of the parameters of the neural network relatively low. The ReLU activation is a function that is linear for inputs > 0 and 0 for inputs ⩽ 0. This type of activation often generate sparse tensors in the hidden layers of the model, reducing the effective number of elements that contribute to the final output. The main advantage of using a RNN is that it can take into consideration the full length protein at once, giving a more accurate and comprehensive prediction. The bidirectional approach ensures that one sequence direction is not *preferred* over the other when analyzing the protein.

**Figure 1:**
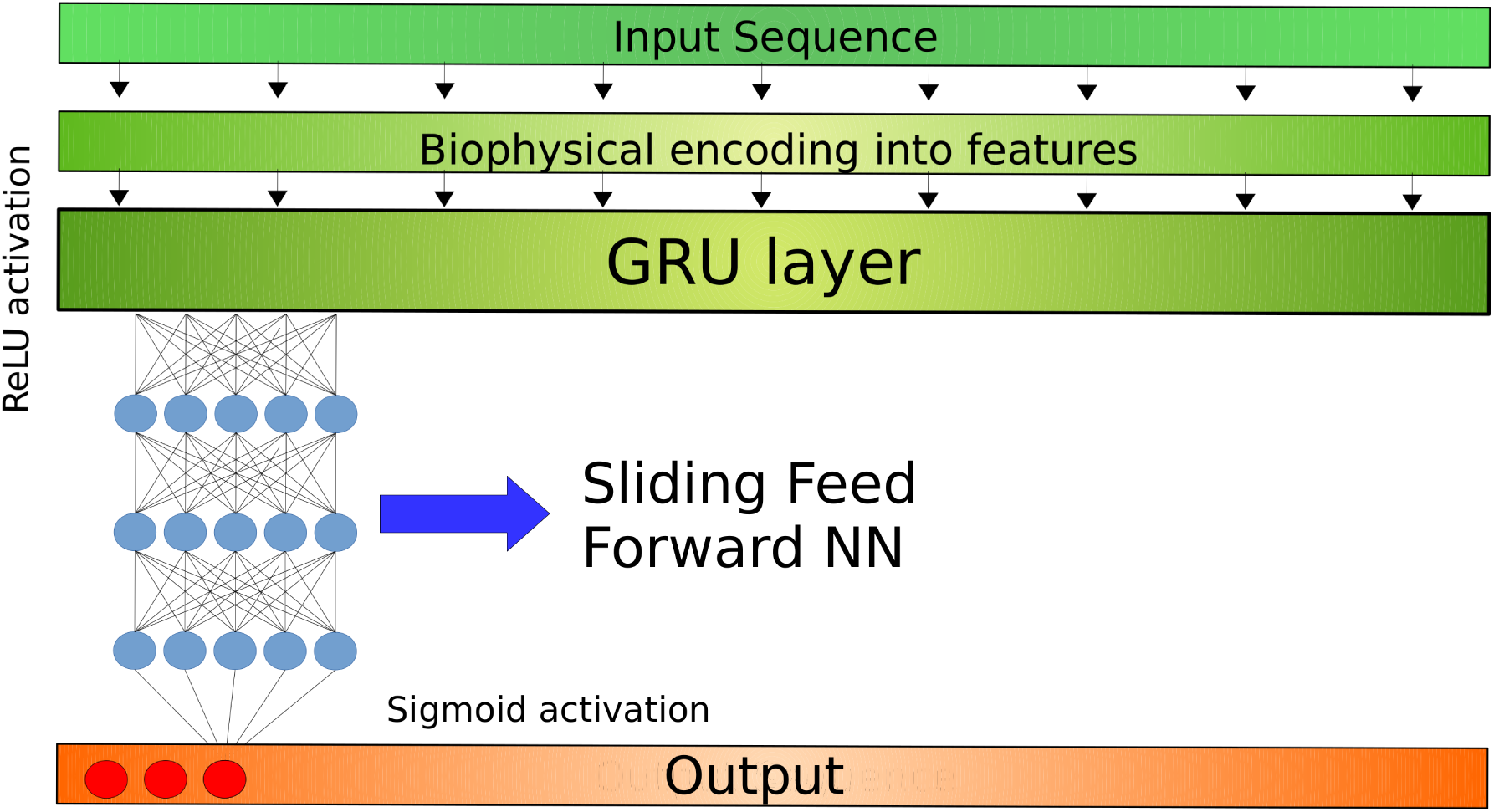
Structure of the neural network: each protein sequence is first translated into feature vectors with shape L × 42, where L is its length. The first part of the NN consists in a Bidirectional Recurrent NN which takes the whole encoded protein as input. The second part of the model is made of a sliding feed forward Neural network (FFNN) with 3 layers of 30 neurons each and ReLU activation. The output layer has sigmoid activation and provides a probability-like final prediction disorder score.

To reduce the dimensionality of the output of the GRU layer, so reducing the parameters of the models, we applied a max-pooling to its output to obtain a tensor of length *L*. In this case maxpooling is used to sample the sequential information provided by the recurrent layer, so limiting overfitting. The output of the max-pooling is passed to a Feed Forward Neural Network (FFNN) sub module using a 11-neurons sliding window to leverage the importance of the local context. The sub module consists of three fully connected layers of 30 neurons each, with ReLU activation. The final neuron has sigmoid activation and it assigns the estimated disorder probability to each amino acid residue. The final network has 17371 parameters that are trained using the Adam algorithm [21] with learning rate of 0.001. The size of the layers has been chosen so that the total number of parameters is about one tenth of the total training data points, this to avoid overfitting.

### 2.4 Evaluation of the results

To evaluate the performances of our method, we reproduced the validation performed in [11], in which 24 predictors were benchmarked on the assessment dataset. The prediction performance was evaluated using the standard Accuracy (ACC), Sensitivity (SEN), Specificity (SPE), Area Under the ROC curve (AUC) and the Matthews Correlation Coefficient (MCC) scores. The performances for other predictors are taken from [11].

## 3 Results

### 3.1 DisoMine performs on par with the state-of-the-art

In Table 1 we show the comparison of the DisoMine per-residue performances with the disorder prediction methods benchmarked in [11], restricting the analysis only to single-sequence methods where no evolutionary information is used to compute their predictions. On this assessment dataset, DisoMine performances are 3% to 34% better in terms of AUC (2.5% to 19.0% in absolute terms) and 21% to 64% (8.6% to 31.8% in absolute terms) higher in terms of MCC. This shows that DisoMine performs well compared to available state-of-the-art single-sequence based methods, and that with respect to sensitivity and specificity, it shows more balanced performances.

**Table 1:**
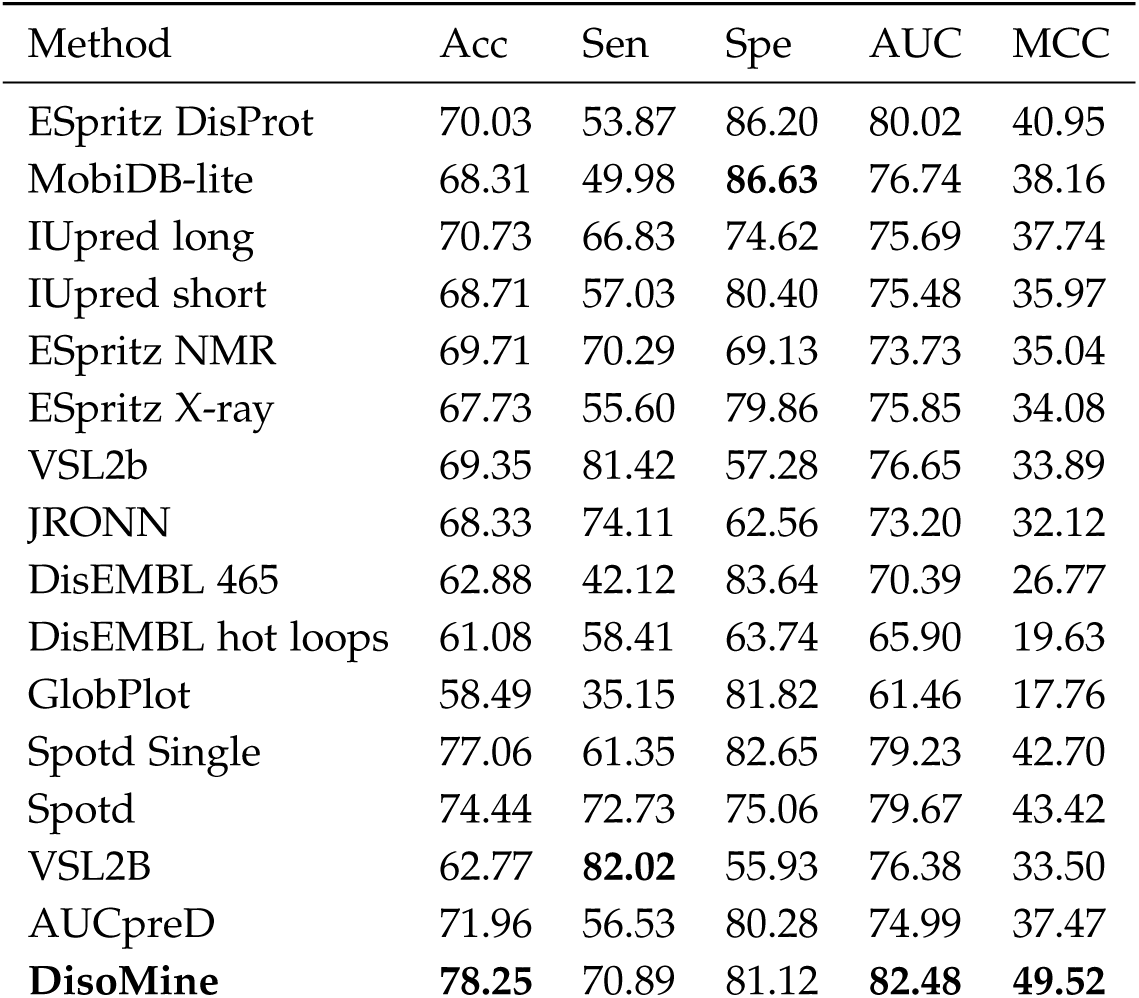
Comparison of the DisoMine performances with state of the art tools. All performances [16, 22, 23, 24, 25, 26, 27, 28, 29, 30], except for Spotd Single, Spotd and VSL2B, are taken from [11]

| Method | Acc | Sen | Spe | AUC | MCC |
| --- | --- | --- | --- | --- | --- |
| ESpritz DisProt | 70.03 | 53.87 | 86.20 | 80.02 | 40.95 |
| MobiDB-lite | 68.31 | 49.98 | **86.63** | 76.74 | 38.16 |
| IUpred long | 70.73 | 66.83 | 74.62 | 75.69 | 37.74 |
| IUpred short | 68.71 | 57.03 | 80.40 | 75.48 | 35.97 |
| ESpritz NMR | 69.71 | 70.29 | 69.13 | 73.73 | 35.04 |
| ESpritz X-ray | 67.73 | 55.60 | 79.86 | 75.85 | 34.08 |
| VSL2b | 69.35 | 81.42 | 57.28 | 76.65 | 33.89 |
| JRONN | 68.33 | 74.11 | 62.56 | 73.20 | 32.12 |
| DisEMBL 465 | 62.88 | 42.12 | 83.64 | 70.39 | 26.77 |
| DisEMBL hot loops | 61.08 | 58.41 | 63.74 | 65.90 | 19.63 |
| GlobPlot | 58.49 | 35.15 | 81.82 | 61.46 | 17.76 |
| Spotd Single | 77.06 | 61.35 | 82.65 | 79.23 | 42.70 |
| Spotd | 74.44 | 72.73 | 75.06 | 79.67 | 43.42 |
| VSL2B | 62.77 | **82.02** | 55.93 | 76.38 | 33.50 |
| AUCpred | 71.96 | 56.53 | 80.28 | 74.99 | 37.47 |
| **DisoMine** | **78.25** | 70.89 | 81.12 | **82.48** | **49.52** |

Table 2 compares DisoMine with the methods tested in [11] that use evolutionary information as part of their pipeline. Evolutionary information is generally collected by computing Multiple Sequence Alignments (MSA) for each protein. These MSAs contain relevant information for many structural bioinformatics problems [31, 17, 32], because at least part of the protein evolutionary history, such as preservation of secondary structure elements, is encoded by the MSA-based features. DisoMine also ranks high in this comparison, providing an AUC improvement of 2.5% and an MCC improvement of 9% with respect to ESpritz X-ray (MSA-based version), the second best method in the benchmark.

**Table 2:** Comparison of the performances of DisoMine with the state of the art tools based on evolutionary information [16, 30, 33, 34, 35, 36, 37, 38], taken from [11].

| Method | Acc | Sen | Spe | AUC | MCC |
| --- | --- | --- | --- | --- | --- |
| ESpritz X-ray | 75.10 | 74.41 | 75.79 | 80.43 | 45.42 |
| ESpritz DisProt | 68.83 | 47.41 | **90.26** | 81.50 | 41.62 |
| AUCpred | 70.39 | 57.66 | 83.11 | 72.77 | 39.98 |
| MetaDisorder | 72.01 | 75.94 | 68.08 | 57.01 | 38.84 |
| MetadisorderMD | 71.56 | 73.78 | 69.34 | 59.52 | 38.22 |
| ESpritz NMR | 71.37 | 75.18 | 67.56 | 76.70 | 37.69 |
| SPINE-D | 71.43 | **81.89** | 60.97 | 78.64 | 37.47 |
| DISOPRED3 | 70.27 | 66.34 | 74.19 | 76.56 | 36.87 |
| MetadisorderMD2 | 70.46 | 77.36 | 63.57 | 69.26 | 35.85 |
| MFDp2 | 68.08 | 57.32 | 78.84 | 67.73 | 34.34 |
| MFDp | 67.18 | 60.50 | 73.86 | 67.40 | 31.50 |
| S2D | 64.15 | 74.05 | 54.25 | 72.11 | 24.79 |
| Metadisorder3D | 51.78 | 43.27 | 60.28 | 61.07 | 3.17 |
| **DisoMine** | **78.25** | 70.89 | 81.12 | **82.48** | **49.52** |

More details about the performance are available for the ROC curve of DisoMine on the test set (Supplementary Figure S1), the relationship between protein length and DisoMine accuracy (Supplementary Figure S2) and the relationship between the length of the predicted disordered region and DisoMine accuracy (Supplementary Figure S3).

We also tested the DisoMine performances on two datasets: the first one is the CASP7 dataset, the second one the PDB dataset described in [16]. We used these two datasets to evaluate its performance on datasets with shorter disorder annotations, as we trained on longer disordered regions from DISPROT (the median disordered regions length is 34). The performances of disomine on the CASP7 and PDB datasets are reported in Supplementary Table 1 and 2 respectively. As expected, the DisoMine performances are lower for this dataset, although they are still meaningful.

### 3.2 DisoMine predicts from a single protein sequence

The excellent overall performance of DisoMine, which only uses single protein sequences as input, has at least two advantages compared to the methods that use evolutionary information. First, it ensures that the quality of the predictions does not depend on the availability of evolutionary information (*i.e.* the number of homologs in the MSAs). This is important, since there is a correlation between the availability of homologous sequences and prediction quality [18]. Similar biases may indeed be present for disorder prediction methods that use evolutionary information, which might so overestimate their prediction performance for the well-studied proteins in the dataset. DisoMine, on the other hand, does not have this limitation and should thus be able to deal with poorly studied and orphan proteins without its prediction performance suffering from the limited evolutionary information available for them. Second, computing MSAs is a time-consuming procedure, and methods that avoid this step are significantly faster, making them amenable to screening entire proteomes. DisoMine can compute the prediction for a protein in about 1-2 seconds on a single core machine, while even the fastest homology detection methods will in our experience typically require minutes even when running in parallel on multiple cores.

### 3.3 Analysis of the predicted patterns

To investigate the patterns DisoMine inferred from the training data, we further analyzed its prediction characteristics on the test set. Figure 2 shows the distribution of the predicted DisoMine scores in relation to the (normalised) position on the sequence. The red line indicates the median, while the two shades of blue indicate percentiles 45-55 and 35-65. The plot shows that one of the patterns learned by DisoMine is that disorder is. as expected, more frequently observed in the C and N terminal regions of proteins, with the overall disorder content gradually decreasing towards the middle of proteins.

**Figure 2:**
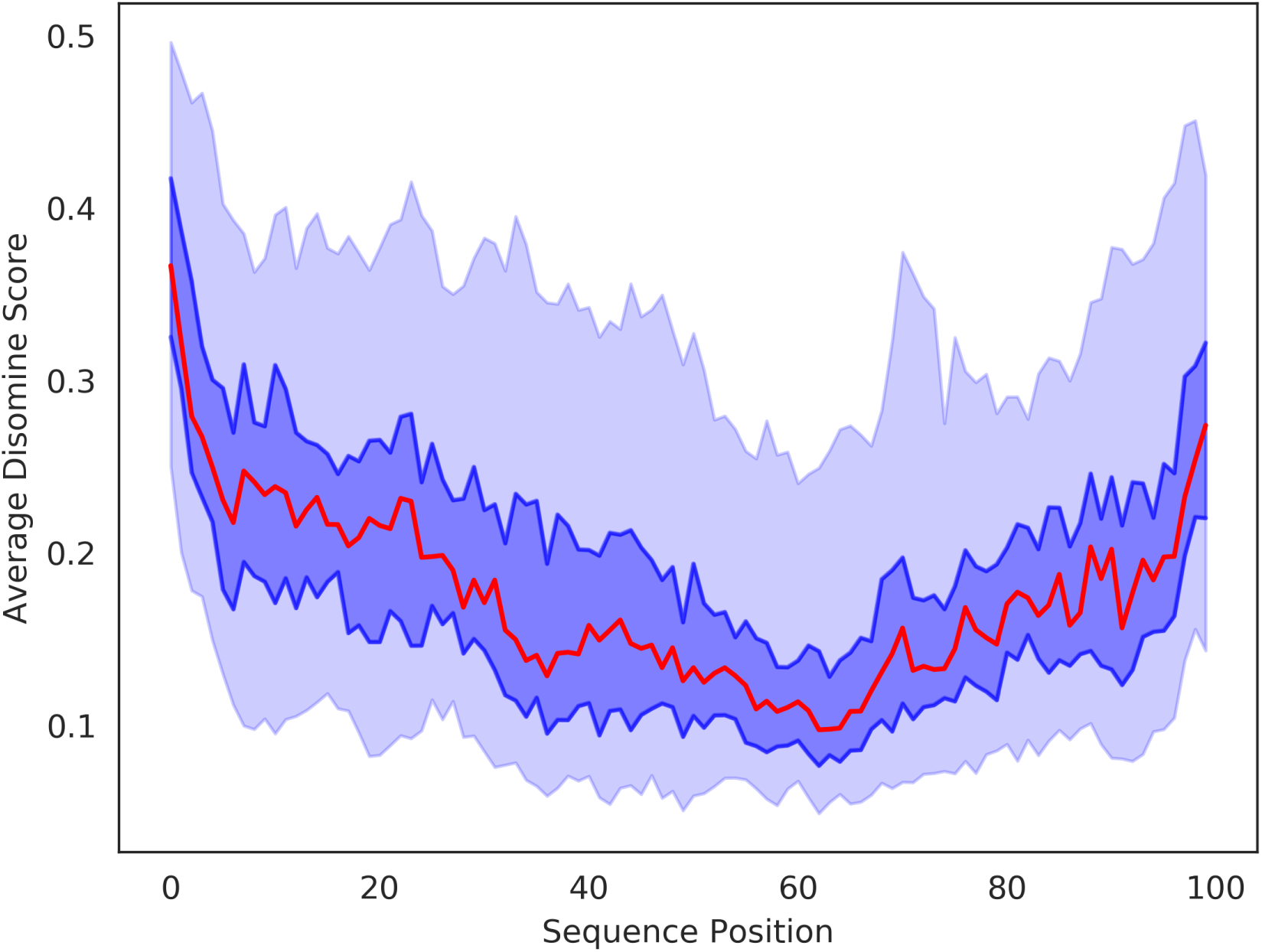
Plot showing the overall distribution of the predicted disorder in relation to protein position. The x-axis represents the position on the protein sequence expressed as fraction of the total length, the y-axis the average DisoMine score. The red line represents the median value. The dark and light blue areas represent respectively the 45-55 and 35-65 percentiles.

### 3.4 Exploring the disorder composition of the Dark proteome

Since many disorder predictors exist, it is not trivial which one to choose for a bioinformatics analysis. To illustrate how this can influence the results of the analysis and its biological conclusions, we reproduced a previous study where we investigated how the performance of structural bioinformatics methods varies with the amount of available evolutionary information [18]. This study highlighted an observation selection bias in the validation process of many protein structure prediction benchmarks, with proteins for which experimentally determined 3D structures are available tending to have significantly more known homologous sequences compared to proteins for which no 3D structure is known. This analysis was based on two sequence datasets: one for 5000 proteins that have a structure in the PDB (STRUCT), another one for 5000 comparable proteins without any such public structure data (NOSTRUCT). In our study we also investigated the role of protein disorder, where we expected to find a weak but significant inverse correlation between i) the percentage of disorder in each protein and ii) the number of homologs available for it. The underlying reason for this would be that alignment tools such as HMMer [39] or Hhblits [40] would have more difficulty in aligning disordered regions [41], which tend to have high sequence variation, and that this would then decrease the number of homologous sequences these methods can find. We did, however, find no such correlation based on IUpred [23] to determine disorder content, and concluded that this link did not exist.

Using DisoMine (Figure 3), however, we do find a correlation: whereas the Spearman correlation with IUpred was -0.087, it is -0.333 with DisoMine. This leads to a different and more straightforward conclusion: there are typically less known homologs for disordered proteins because their inherent higher sequence variability makes them more difficult to align. This difference between the predicted disorder is also evident from their distribution, here expressed as the mean of the predictions over all residues per sequence in the STRUCT and NOSTRUCT datasets (Fig. 4). The median predicted disorder content is respectively 0.187 and 0.337 for Disomine, and 0.247 and 0.302 for IUPred. Although both tools indicate that proteins without known structure tend to have more disordered residues, the difference between the two DisoMine predicted distributions is larger, with the shape of the distribution also visibly different. This illustrates how the two methods identify disorder in distinct ways, and highlights how the choice of bioinformatics software influences the (biological) conclusions that are drawn. It is likely best to at least explore several different ones when performing such analyses, and what the predictor is best at. In this case, for example, IUPRED likely works much better on short disordered regions, but such regions will not influence the overall sequence alignment methods much, whereas longer disordered regions as picked up by DisoMine will.

**Figure 3:**
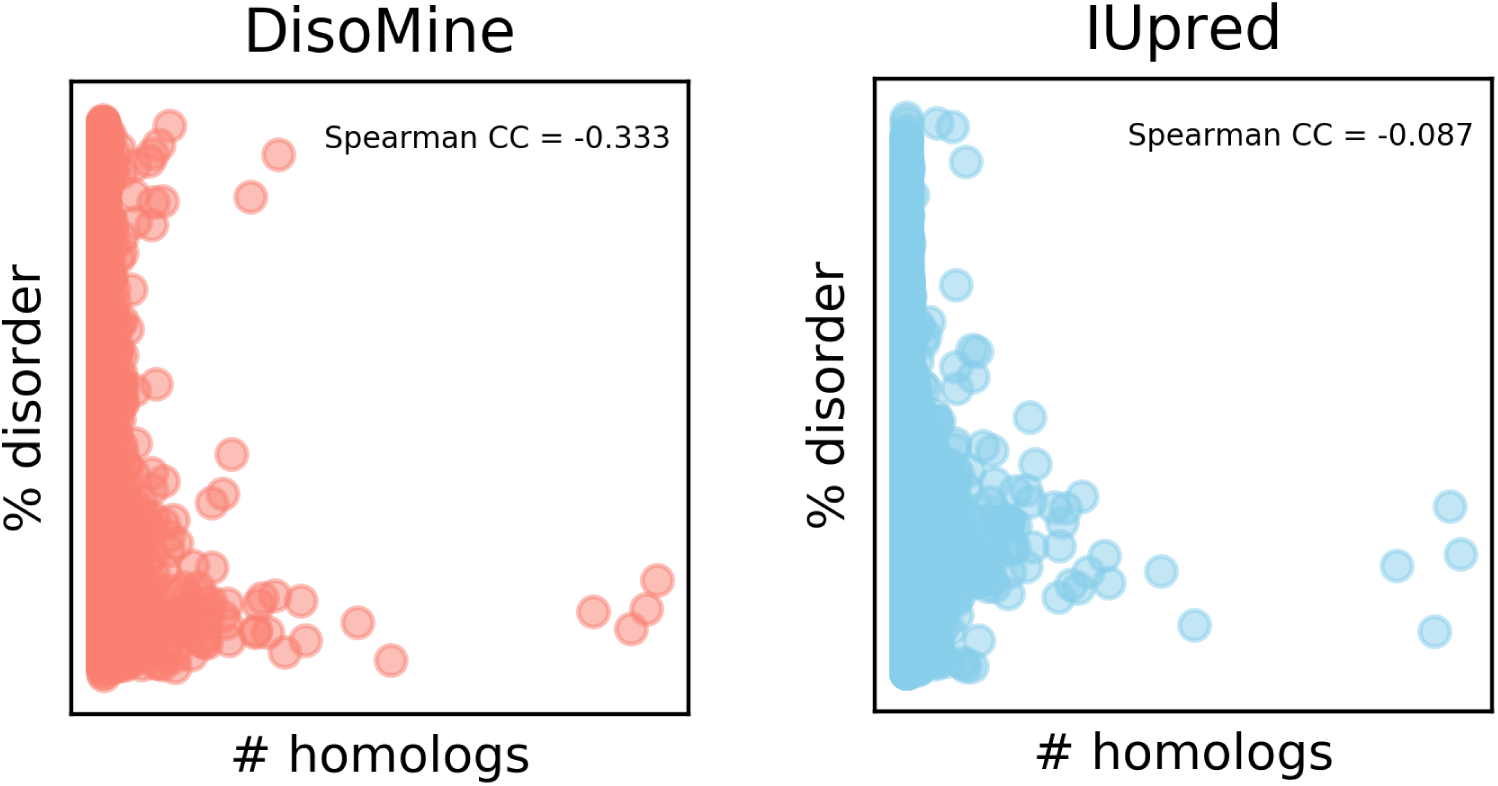
Correlation between disorder content and number of available homologs for DisoMine (top) and IUpred (bottom).

**Figure 4:**
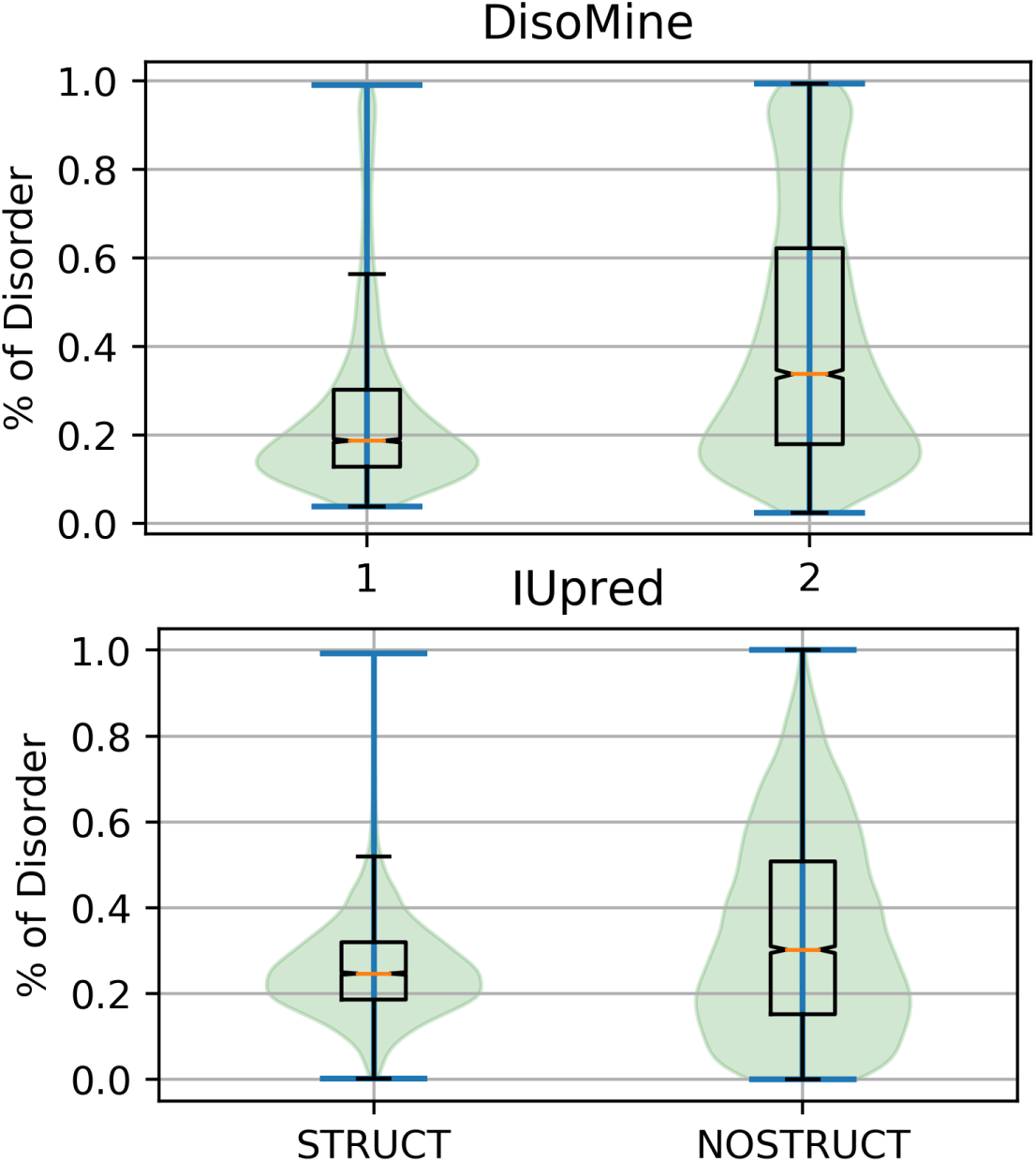
Distributions of the predicted disorder content, as percentage of disordered residues in the protein, as predicted by DisoMine (top) and IUpred (bottom) for the STRUCT and the NOSTRUCT datasets.

## 4 Webserver availability

DisoMine is available via the http://bio2byte.com/disomine/ web server. Only the target protein sequences in the FASTA format are required as input. The webserver can be accessed either via the browser or programmatically via a REST API, and provides residue level predictions of disorder within seconds. The results are downloadable as text or can be interactively visualized via a JavaScript applet. The web server can also provide EFoldMine and DynaMine predictions.

## 5 Conclusion

DisoMine predicts protein disorder from single protein sequences in seconds, while performing on par with current state of the art methods on a publicly available large-scale benchmark. It does so using predicted biophysical characteristics, not the protein amino acid sequence directly, and might so provide more generally applicable disorder predictions. Based on the DisoMine approach, we show that the choice of protein disorder predictor can strongly influence a bioinformatics analysis. This can potentially lead to different biological hypotheses, and might lead to interpretation errors that can be propagated in scientific literature. Adopting the best predictors available, while taking care that they are the most relevant for the problem at hand (e.g. long vs. short disorder), then ensuring several predictors are used in parallel should alleviate these issues. DisoMine is available as web server, with an user-friendly interface and API acces that allows automatic interaction with the server for large scale analysis and scripting.

## Funding

GO acknowledges funding by the Research Foundation Flanders (FWO) project nr. G.0328.16N. DR is funded by a KU Leuven post-doctoral mandate and an FWO post-doctoral fellowship. This work was supported by the European Regional Development Fund (ERDF) and the Brussels-Capital Region-Innoviris within the framework of the Operational Programme 2014-2020 through the ERDF-2020 project ICITY-RDI.BRU. Part of the work has been developed thanks to the COST Action BM1405 NGP-net.

## Acknowledgements

G. O. and D. R. are grateful to A. Motz, A. L. Mascagni for the support and the helpful discussions, and to R. Gilbert and R. Sanchez for the inspiration.

## Notes

http://bio2byte.com/disomine/

### Competing Interest Statement

The authors have declared no competing interest.

