## Supplementary for "Prediction of disordered regions in proteins with recurrent Neural Networks and protein dynamics"

### Supplemantary Material

May 25, 2020

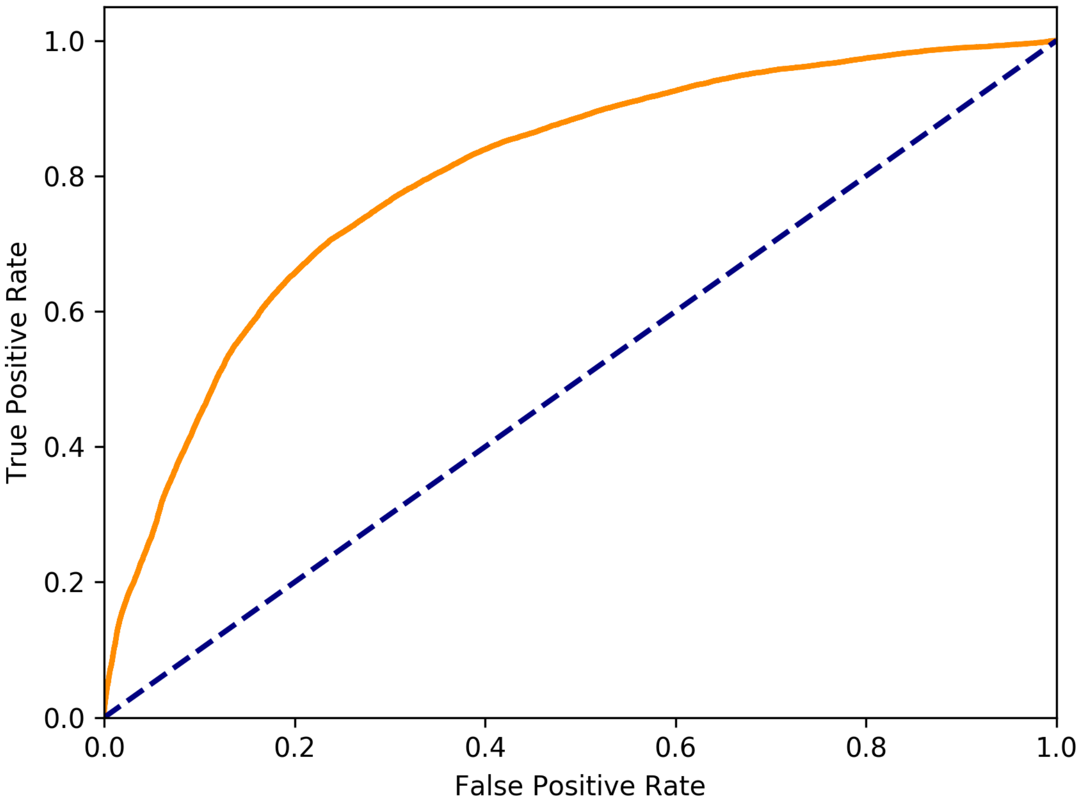


Figure S1: Roc curve of DisoMine, tested on the DisProt test dataset

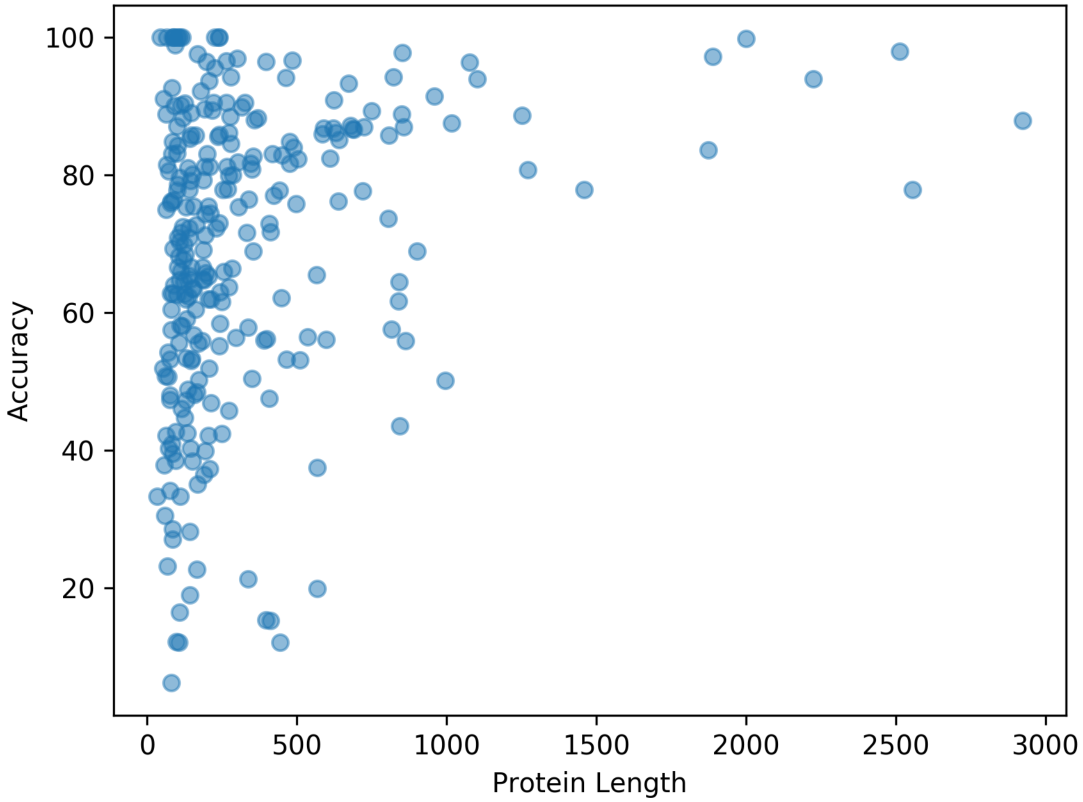


Figure S2: Scatter plot that relates the length of the predicted disordered region with DisoMine accuracy. Every blue dot represents a protein region predicted as disordered

Table S1: Table showing the performances of DisoMine and other predictors on the CASP7 dataset

| Method | Acc | Sen | Spe | AUC | MCC |
|---|---|---|---|---|---|
| vsl2b | 54.05 | 83.77 | 52.16 | 81.17 | 17.06 |
| SPOT-D | 55.18 | 93.52 | 52.74 | 87.16 | 21.96 |
| AUCPred | 54.33 | 81.9 | 52.77 | 79.4 | 15.6 |
| SPOT-D single | 54.4 | 86.04 | 52.39 | 82.72 | 18.23 |
| DisoMine | 54.07 | 83.93 | 52.17 | 76.58 | 17.15 |

Table S2: Table showing the performances of DisoMine on the PDB dataset

| Method | Acc | Sen | Spe | AUC | MCC |
|---|---|---|---|---|---|
| DisoMine | 68.17 | 78.74 | 67.44 | 78.14 | 23.61 |

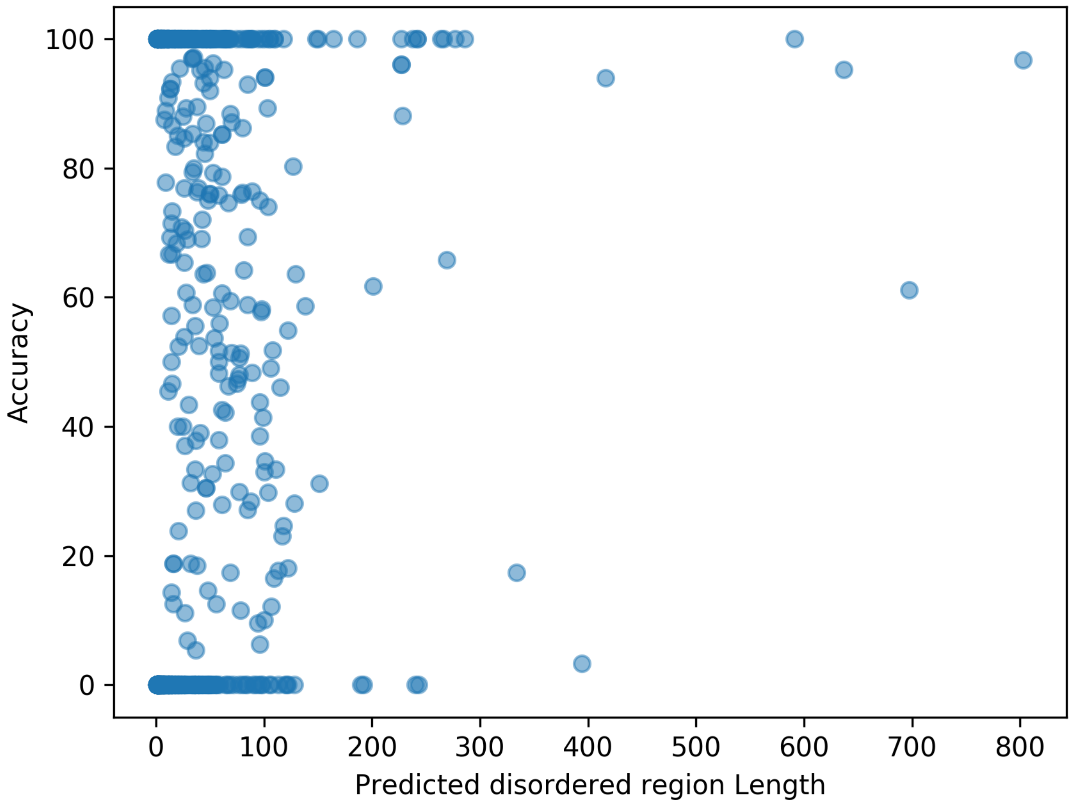


Figure S3: Scatter plot that relates protein length with DisoMine accuracy. Every blue dot represents a protein and the plot shows the relationship between protein length and the accuracy on that specific protein

#### S1 STRUCT and NOSTRUCT datasets

The dataset of proteins associated with solved structures have been selected from the PDB [?] database, taking the structures with resolution lower than 2Å. We then applied sequence filtering: first of all we removed all the proteins shorter than 50 or longer than 1500 amino acids. Secondly we removed sequence redundancy using BLASTCLUST [?] with a maximum sequence identity of 20% and coverage of 90%. From the remaining sequences, we sampled 5000 proteins. We called them the STRUCT dataset and they are supposed to be a representative sample of poly-peptides with solved crystal structure. The dataset of proteins with no solved structures have been built starting from the Uniprot sequences with existence evidence at the transcriptomic or proteomic level. We ran a BLAST search against PDB in order to select the sequences that had no evolutionary relationship with any of the proteins with solved structure. We applied the same sequence filtering and again we sampled 5000 protein from the remaining proteins. This dataset is meant to be representative of the proteins without solved structures and we called it NOSTRUCT.For more details about this analysis see [?].
